# Therapeutic effects of Hypoxia-Inducible Factor-1α (HIF-1α) on bone formation around implants in diabetic mice

**DOI:** 10.1101/392670

**Authors:** Sang-Min Oh, Jin-Su Shin, Il-Koo Kim, Jae-Seung Moon, Jung-Ho Kim, Sang-Kyou Lee, Jae-Hoon Lee

## Abstract

Patients with uncontrolled diabetes are susceptible to implant failure due to impaired bone metabolism. Hypoxia-Inducible Factor 1α (HIF-1α), a transcription factor that is up-regulated in response to reduced oxygen condition during the bone repair process after fracture or osteotomy, is known to mediate angiogenesis and osteogenesis. However, its function is inhibited under hyperglycemic conditions in diabetic patients. The aim of this study is to evaluate the effects of exogenous HIF-1α on bone formation around implants by applying HIF-1α to diabetic mice via a novel PTD-mediated DNA delivery system. Smooth surface implants (1mm in diameter; 2mm in length) were placed in the both femurs of diabetic and normal mice. HIF-1α and placebo gels were injected to implant sites of the right and left femurs, respectively: Normal mouse with HIF-1α gel (NH), Normal mouse with placebo gel (NP), Diabetic mouse with HIF-1α gel (DH), and Diabetic mouse with placebo gel (DP). RNA sequencing was performed 4 days after surgery. Based on RNA sequencing, Differentially Expressed Genes (DEGs) were identified and HIF-1α target genes were selected. Histologic and histomorphometric results were evaluated 2 weeks after the surgery. The results showed that bone-to-implant contact (BIC) and bone volume (BV) were significantly greater in the DH group than the DP group (p < 0.05). A total of 216 genes were differentially expressed in DH group compared to DP group. On the other hand, there were 95 DEGs in the case of normal mice. Twenty-one target genes of HIF-1α were identified in diabetic mice through bioinformatic analysis of DEGs. Among the target genes, *NOS2, GPNMB, CCL2, CCL5, CXCL16* and *TRIM63* were manually found to be associated with wound healing-related genes. In conclusion, local administration of HIF-1α via PTD may help bone formation around the implant and induce gene expression more favorable to bone formation in diabetic mice.

## Introduction

Dental implant has become an efficient and predictable treatment for replacing missing teeth. The number of implants placed in the United States has been steadily increasing at 12#x0025; annually, with improvements in implant materials, designs, and surgical techniques [1]. Despite its high success rate of 95% in the general population [2], certain risk factors may predispose individuals to implant failure [3]. Among various patient-related risk factors, poorly controlled diabetes mellitus, a chronic metabolic disease characterized by hyperglycemia, has been considered a relative contraindication to dental implant [4–6].

Implant success is highly dependent on osseointegration, a process in which bone and implant surface become structurally and functionally integrated without interposition of non-bone tissue layer [7]. Osseointegration involves bone repair and remodeling and critically affects implant stability [8]. However, the hyperglycemic condition of diabetes inhibits osteoblastic differentiation, mineralization, and adherence of extra cellular matrix and stimulates bone resorption, consequently interfering with wound healing and bone regeneration [9,10]. Previous experimental studies have reported decreased bone-to-implant contact (BIC) and delayed new bone formation around the implant in diabetic animal models, proving that hyperglycemia impairs osseointegration [11].

Numerous studies have demonstrated that chronic high glucose level results in defective responses of tissues to hypoxic conditions by impairing the function of Hypoxia-inducible factor 1α (HIF-1α) [12–14]. Transcription factor HIF-1α is up-regulated in response to reduced oxygen conditions and influences numerous target genes such as vascular endothelial growth factor (VEGF) and runt-related transcription factor 2 (RUNX2), which are known to be associated with angiogenesis and osteogenesis [15–17]. HIF-1α, which is well known to play a pivotal role in wound healing, is stabilized against degradation and transactivates under hypoxia [15].

A study carried out by Zou et al. demonstrated that osteogenesis and angiogenesis were enhanced around implants with the up-regulation of HIF-1α in rat bone mesenchymal stem cells (BMSCs) in animal models. In addition, previous studies investigating effects of HIF-1α on bone regeneration showed that functions of osteoblasts and chondrocytes are directly regulated by HIF-1α during bone fracture healing in animal models [18,19]. As many studies have associated malfunction of HIF-1α in diabetic animal models with delayed bone recovery, attempts have been made to improve bone healing by applying HIF-1α. However, application of HIF-1α using mesenchymal stem cells to increase expression of HIF-1α is limited by its inefficiency and long preparation period. To maximize efficiency of delivery to the implant site, a PTD-mediated DNA delivery system was used in this study. PTDs are short peptides efficiently transporting various proteins, nucleic acids, and nanoparticles into cells across the plasma membranes. This protein-based strategy is a remarkable method for delivering target DNA to the nucleus because of its low toxicity and high transduction efficiency [20]. Indeed, it has been shown that overexpression of HIF-1α induced by PTD-mediated DNA delivery system resulted in increased expression of VEGF and angiogenesis in vitro and in vivo [21].

In this study, taking advantage of the fact that PTD can deliver HIF-1α into cell nuclei, we designed an experiment to determine whether local application of HIF-1α into the implanted sites by using PTD in the femur of diabetic mice enhances osseointegration compared with placebo controls. Using RNA sequencing and histomorphometric analysis, we observed new bone formation and significant changes in the expression of genes associated with wound healing.

## Materials and methods

### Ethics statements

The research proposal was approved by the Laboratory Animal Care and Use Committee at Yonsei University Biomedical Research Institute (Approval number 2014-0032). Animals received an intraperitoneal injection of a mixture of Zoletil 50^®^ (A mixture of zolazepam and tiletamine; Vibac Laboratories, Carros, France; 30mg/kg body weight) and Rompun^®^ (Xylazine hydrochloridine; Bayer, Leverkusen, Germany; 10mg/kg body weight) to minimize suffering and pain. Animals were exposed to CO_2_ gas for euthanasia. The ARRIVE Guidelines for reporting animal research were abided by in all sections of this report [22].

### Animal models

Thirteen 8-week-old male C57BL/6 mice (21 g) from Charles River (Orientbio, Gapyeong-gun, Korea) and thirteen 8-week-old male C57BLKS/J-db/db mice (38 g, Leptin-receptor deficient type 2 diabetes mice) from Charles River (Hinobreeding Center, Tokyo, Japan) were used for the experiments. They were maintained in the Avison Biomedical Research Center at Yonsei University College of Medicine at 23 ± 2°C and 50 ± 10% humidity under 12 hours of light alternating with 12 hours of darkness.

### Preparation of HIF-1α gene construct and Hph-1-GAL4 DNA binding domain protein

HIF-1α encoded plasmid and Hph-1-GAL4 DNA binding protein were provided by Sang-Kyou Lee’s Laboratory at Yonsei University, Department of Biotechnology. The HIF-1α gene was inserted into pEGFP-N1 UAS plasmid containing five consensus GAL4 binding sites (UAS: CGGAGGACAGTACTCCG) (HIF-1α-UAS). The GAL4 DNA binding domain that encodes the DNA-interactive domain of yeast transcription factor GAL4 was cloned into pRSETB plasmid (Clonetech) expression vector containing Hph-1-PTD sequence (YARVRRRGPRP) at the N-terminus (Hph-1-GAL4-DBD). pRSETB plasmid with the Hph-1-GAL4 DNA binding domain was transformed into *Escherichia coli* BL21 star (DE3) pLysS strain (Invitrogen). Protein expression and purification were performed as described previously [20].

### Preparation of HIF-1α gel

One microgram of HIF-1α-UAS plasmid was mixed with 50 μg of Hph-1-GAL4-DBD at room temperature for 15 min right before surgery, as previously described [21]. The liquid form of Matrigel^®^ (BD Biosciences, San Jose, CA, USA) and the mixture were blended at a 1:1 ratio just before the application of HIF-1α gel during surgery (Fig 1). Pure Matrigel^®^ was used as a placebo gel. Matrigel^®^ was stored in a liquid state at temperature −72°C in the freezer because it solidifies at 4°C.

**Fig 1.**
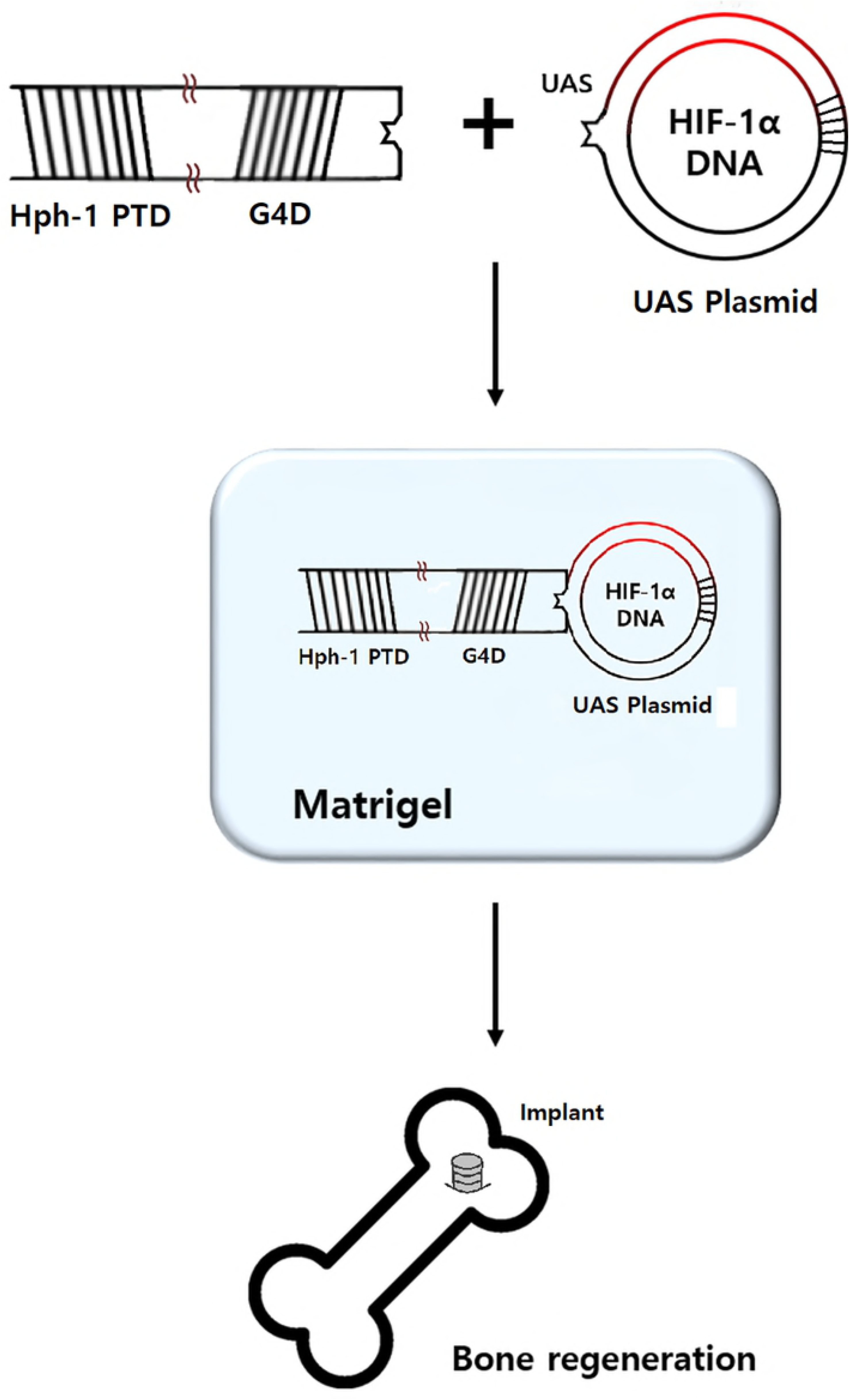
Preparaion of HIF-1α gel. GAL4 DNA binding domain (G4D) was cloned into pRSETB plasmid expression vector containing Hph-1-PTD sequence at the N-terminus (Hph-1-G4D). The HIF-1α gene was inserted into pEGFP-N1 UAS plasmid containing five consensus GAL4 binding sites (HIF-1α-UAS plasmid). HIF-1α-UAS plasmid and Hph-1-G4D were mixed at a 1:50 mass ratio. The mixture and liquid form of Matrigel^®^ were blended at a 1:1 ratio just before application for bone regeneration.

### Surgical procedure

Thirteen C57BL/6 mice (21 g) and thirteen C57BLKS/J-db/db mice (38 g) were given two weeks of acclimatization before surgery. Implant placing methods followed Xu et al. (Xu *et al.*, 2009). The mice were anesthetized by intraperitoneal injection of a mixture of Zoletil 50^®^ (30 mg/kg) and Rompun^®^ (10 mg/kg) (Fig 2A), the surgical site being shaved (Fig 2B) and disinfected with Betadine. An incision was made above both knee joints and the anterior-distal aspect of the femur was accessed using medial parapatellar arthrotomy (Fig 2C). Implant sites were prepared on the anterior-distal surface of the femur through sequential drilling with 0.5 mm, 0.9 mm round bur and 0.7 mm stainless steel twist drills with cooled sterile saline irrigation (Fig 2D). To effectively deliver HIF-1α to the implant site via local injection, gel phase materials were prepared as described previously. HIF-1α gel was injected to the preparation site and cancellous bone of the right femur, and placebo gel was injected to the same areas for the left femur (Fig 2E). When the gel hardened, pure titanium implants with a machined surface (1 mm in diameter; 2 mm in length; Shinhung, Seoul, Korea) were inserted into the undersized hole with mild pressure (Fig 2F). The muscles and skin were sutured independently to cover and stabilize the implant (Fig 2G and Fig 2H). Antibiotics were injected at fixed times daily for 3 days (Enrofloxacin 5mg/kg, twice a day; Meloxicam, 1mg/kg, once a day) [23,24]. Three C57BL/6 (21 g) and three C57BLKS/J-db/db (38 g) mice were sacrificed 4 days after the surgery for RNA sequencing, and ten mice from each strain were sacrificed two weeks after the surgery for histologic and histomorphometric analysis.

**Fig 2.**
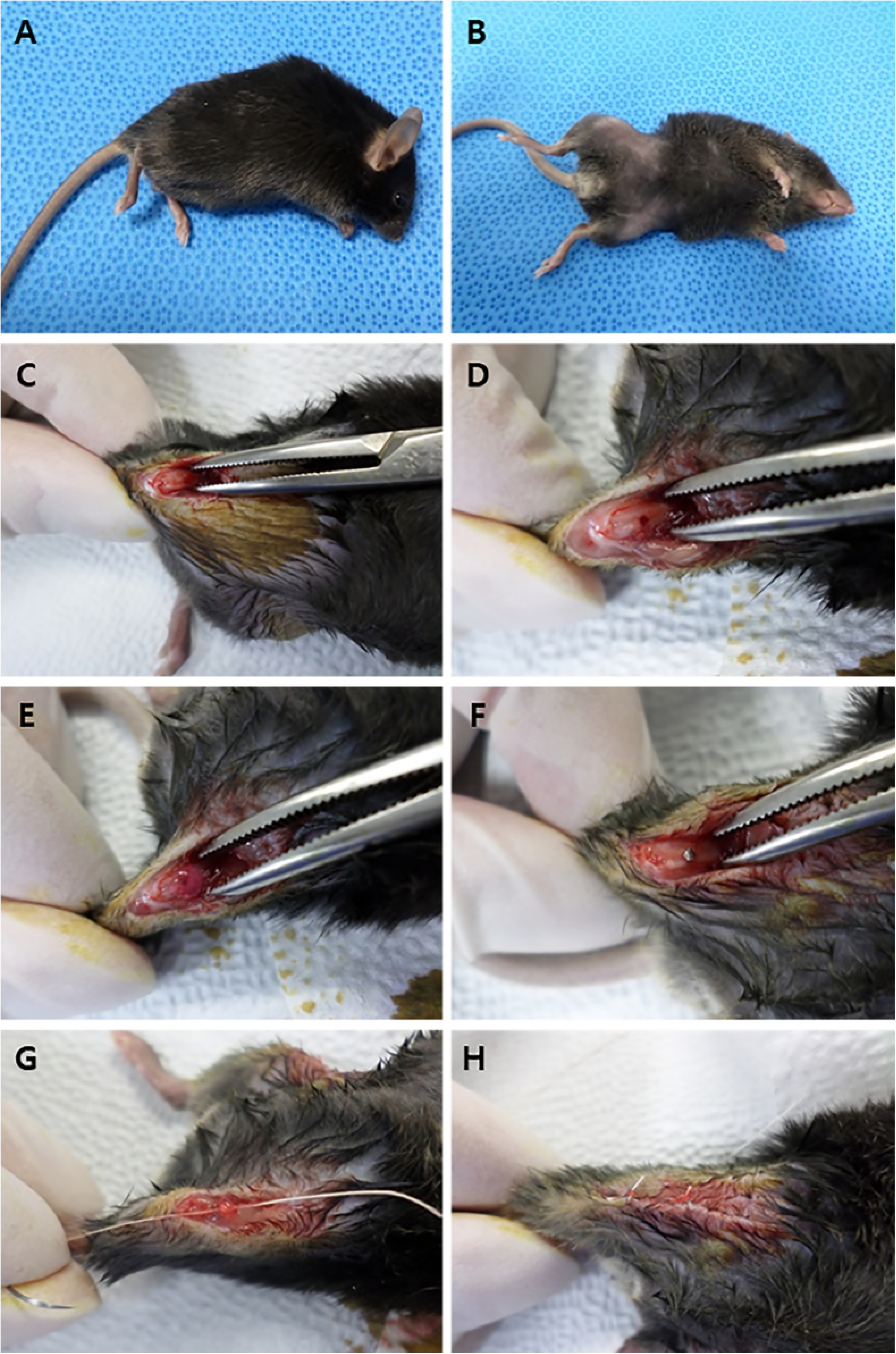
Surgical procedure of implant. (A) Anesthetized mouse, (B) Skin preparation, (C) Incision, (D) Preparation of implant site, (E) Gel injection, (F) Placement of implant, (G) Closure of surgical site layer by layer, (h) Post surgery.

### Measurement of blood glucose levels

1 mm tail cut was obtained from conscious mice, and blood glucose levels from caudal artery were measured twice, before the surgery and sacrifice, using a blood glucose meter and strips (Accu-Check Performa, Roche Diagnostics). The mice were fasted for 12 hours before the measurement [25].

### RNA sequencing

Three mice from each strain (C57BL/6 (21 g) and C57BLKS/J-db/db (38 g)) were sacrificed for RNA sequencing analysis 4 days after implantation. Classification of groups and samples are described in Table 1. Group NH, Normal mice with HIF-1α gel; group NP consisted of Normal mice with placebo gel; group DH, Diabetic mice with HIF-1α; group DP, Diabetic mice with placebo gel. Different combinations of groups were designed and RNA sequencing was performed to identify DEGs.

**Table 1.**
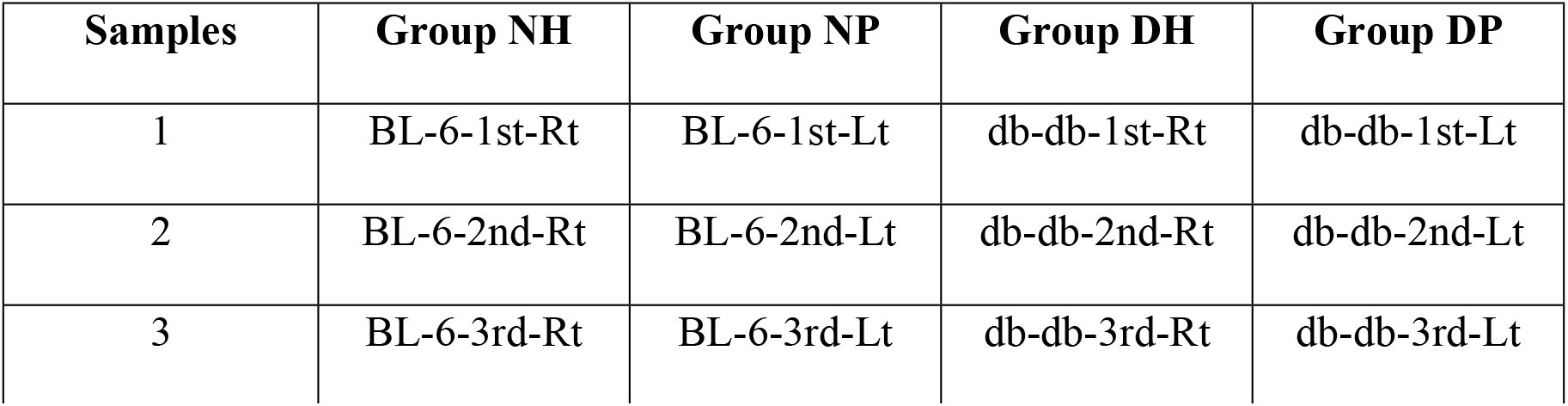
Experimental design for RNA sequencing‥.

RNA purity was determined by assaying 1 μl of total RNA extract on a NanoDrop8000 spectrophotometer. Total RNA integrity was checked using an Agilent Technologies 2100 Bioanalyzer with an RNA Integrity Number (RIN) value greater than 8. mRNA sequencing libraries were prepared according to the manufacturer’s instructions (Illumina TruSeq RNA Prep Kit v2). mRNA was purified and fragmented from total RNA (1 ug) using poly-T oligo-attached magnetic beads using two rounds of purification. Cleaved RNA fragments primed with random hexamers were reverse transcribed into first strand cDNA using reverse transcriptase and random primers. The RNA template was removed and a replacement strand was synthesized to generate double-stranded (ds) cDNA. End repair, A-tailing, adaptor ligation, cDNA template purification and enrichment of the purified cDNA templates using PCR were then performed. The quality of the amplified libraries was verified by capillary electrophoresis (Bioanalyzer, Agilent). After performing QPCR using SYBR Green PCR Master Mix (Applied Biosystems), we combined libraries that were index tagged in equimolar amounts in the pool. RNA sequencing was performed using the Illumina NextSeq 500 system following the protocols provided for 2 x 75 sequencing.

Reads for each sample were mapped to the reference genome (mouse mm10) by Tophat (v2.0.13). The aligned results were added to Cuffdiff (v2.2.0) to report differentially expressed genes. Geometric and pooled methods were applied for library normalization and dispersion estimation.

Each group has three samples. Normal mice with HIF-1α gel (NH); Normal mice with placebo gel (NP); Diabetic mice with HIF-1α gel (DH); Diabetic mice with placebo gel (DP).

### Identification of DEGs

Of the various Cuffdiff output files, “gene_exp.diff’ was used to identify differentially expressed genes (DEGs). Two filtering processes were applied to detect DEGs between control and case groups. First, only genes having Cuffdiff status code “OK” were extracted. The status code indicates whether each condition contains enough reads in a locus for a reliable calculation of expression level, “OK” indicating that the test was successful in calculating gene expression level. For the second filtering, 2-fold change was calculated and only genes belonging to the following range were selected:

Up-regulated: log_2_[case]-log_2_[control] > log2(2) = 1
Down-regulated: log_2_[case]-log_2_[control] < log2(1/2) = −1

### Identification of target genes of HIF-1α

The software program TRANSFAC^®^ (Qiagen N.V., USA) was used to select target genes of HIF-1α. TRANSFAC^®^ provides not only a database of eukaryotic transcription factors but also an analysis of transcription factor binding sites. MATCH analysis was performed with TRANSFAC^®^ using DEGs and an HIF-1α related matrix was selected from the results [26].

### Histologic and histomorphometric analysis

Ten mice from each strain (C57BL/6 (21 g) and C57BLKS/J-db/db (38 g)) were sacrificed at 2 weeks after implant surgery. Femurs from both sides were obtained and fixed at 10% buffered formalin. After a week of fixation, they were embedded in light curing epoxy resin. The specimens were prepared with a cutting distance of 0.5 mm from the apical end of the implant. Sections were cut with a thickness of 50 um via grinding system, stained with hematoxylin and eosin (H&E), and observed using light microscopy (Leica DM LB, Germany). IMT iSolution Lite ver 8.1^®^ (IMT i-Solution Inc., Canada) was used for histomorphometric measurement. BIC ratio was calculated as the linear percentage of direct BIC to the total surface of implants. BV ratio was calculated as the percentage of newly formed bone area to a circumferential zone within 100 um of the implant surface.

### Statistical analysis

All statistical procedures were performed using IBM SPSS 23.0 (IBM Corp., Armonk, NY, USA). Raw histomorphometric measurement data were used to calculate the mean ± SD. Shapiro-Wilk test was used to test normality and one-way analysis of variance (ANOVA) was used to compare groups, which were considered independent. Post hoc was performed with Scheffe’s method. The value of P < 0.05 was considered statistically significant.

## Results

### Blood glucose level

The average glucose level in normal mice before the surgery and sacrifice was 129.6 (mg/dL) and 129.8 (mg/dL), respectively, while it was 477.9 (mg/dL) and 456.1 (mg/dL) in diabetic mice (Table 2). The mice were diagnosed as diabetic if the blood glucose level was over 250 (mg/dL). The blood glucose levels were significantly higher in diabetic mice, much greater than 250 (mg/dL), while those in normal mice were less than 150 (mg/dL) [27,28].

**Table 2.** Blood glucose levels (means ± SDs) of diabetic and normal mice.

|  | Diabetic mice |  | Normal mice |  |
| --- | --- | --- | --- | --- |
| Blood glucose<br>(mg/dL) | Before surgery<br>(n=10) | Before sacrifice<br>(n=10) | Before surgery<br>(n=10) | Before sacrifice<br>(n=10) |
| | 477.9 ± 29.7 | 456.1 ± 51.0 | 129.6 ± 9.5 | 129.8 ± 8.0 |

### RNA sequencing and DEGs

The number of up- and down-regulated genes with a certain cutoff (2-fold; p-val < 0.05; FDR < 0.1) for all combinations are described in Table 3. A total of 216 genes were differentially expressed in the DH group compared to the DP group. On the other hand, there were 95 DEGs in the case of normal mice.

**Table 3.**
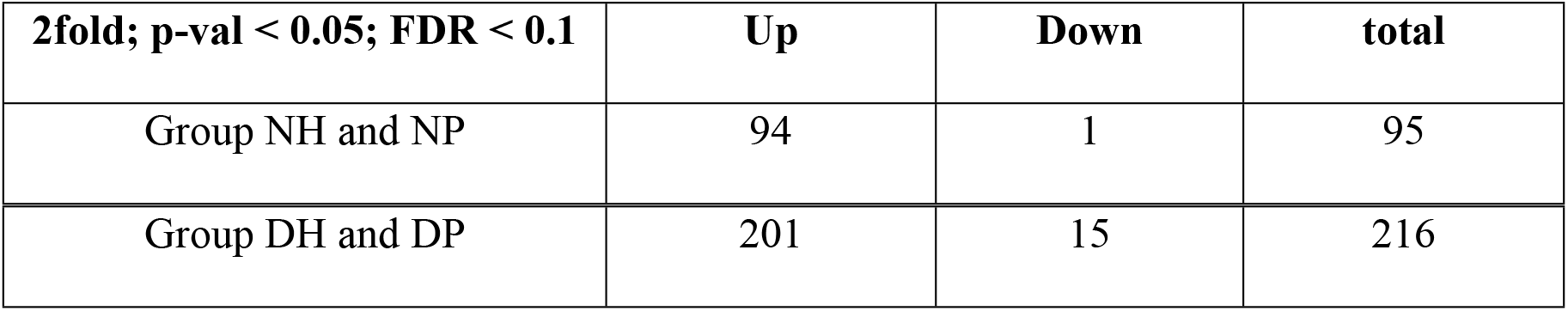
The number of differentially expressed genes (DEG) in each combination.

| 2fold; p-val < 0.05; FDR < 0.1 | Up | Down | total |
| --- | --- | --- | --- |
| Group NH and NP | 94 | 1 | 95 |
| Group DH and DP | 201 | 15 | 216 |

### Target genes of HIF-1α in bioinformatic analysis

TRANSFAC^®^ analysis was performed among DEGs in order to find genes previously reported as targets of HIF-1α. Twenty-one genes were identified as target genes of HIF-1α in diabetic mice (Table 4). Among the 21 detected genes, *NOS2, GPNMB, CCL2, CCL5, CXCL16* and *TRIM63* were found to be associated with wound healing or bone healing-related genes. In normal mice, five genes (*NOS2, CCL2, CCL5, CD274, TNF*) were identified as target genes of HIF-1α.

**Table 4.** 21 target genes of HIF-1α out of 216 differentially expressed genes in diabetic mice.

| Gene symbol | Fold change (log2X)<br>in diabetic mice | Molecule type |
| --- | --- | --- |
| CACNA1S | 1.07 | Calcium channel |
| <u>CCL2</u> | 1.31 | Chemokine |
| <u>CCL5</u> | 1.49 | Chemokine |
| CD274 | 1.70 | Ligand |
| <u>CXCL16</u> | 1.11 | Chemokine |
| COBL | 1.46 | Cordon bleu |
| DES | 1.20 | Enzyme |
| <u>GPNMB</u> | 1.06 | ECM |
| IL2RA | 1.76 | Binding protein |
| JSRP1 | 1.05 | Membrane protein |
| MARCO | 2.88 | Binding protein |
| MIA | -1.03 | Structural protein |
| MURC | 1.12 | Structural protein |
| MYH14 | 1.14 | Enzyme |
| MYL3 | -1.54 | Structural protein |
| <u>NOS2</u> | 1.41 | Enzyme |
| OAS2 | 1.21 | Enzyme |
| PRKAG3 | 1.50 | Protein kinase |
| TGTP1 | 2.16 | GTPase |
| <u>TRIM63</u> | 1.93 | Ubiquitin protein ligase |
| TTN | 1.05 | ECM |

### Histologic analysis

In the NH group, abundant and smooth-lined mature bone formation was observed (Fig 3A and 3B). Mature and smooth-lined bone was also observed in the NP group, but in a lesser amount than in the NH group (Fig 3C and 3D). In contrast, most of the implant surface in the DP group showed soft tissue attachment and abundant adipose tissue in surrounding areas (Fig 3E and 3F). Moreover, bone formation was irregular. In the DH group, many vascular sinusoids with red blood cells (RBC) were located around implants, and more bone formation and attachment around implant were observed than in the DP group (Fig 3G and 3H). In addition, there was a tendency toward increased bone formation at the HIF-1α application site around implants. Bone marrow was filled with adipose tissue in areas distant from the application site.

**Fig 3.**
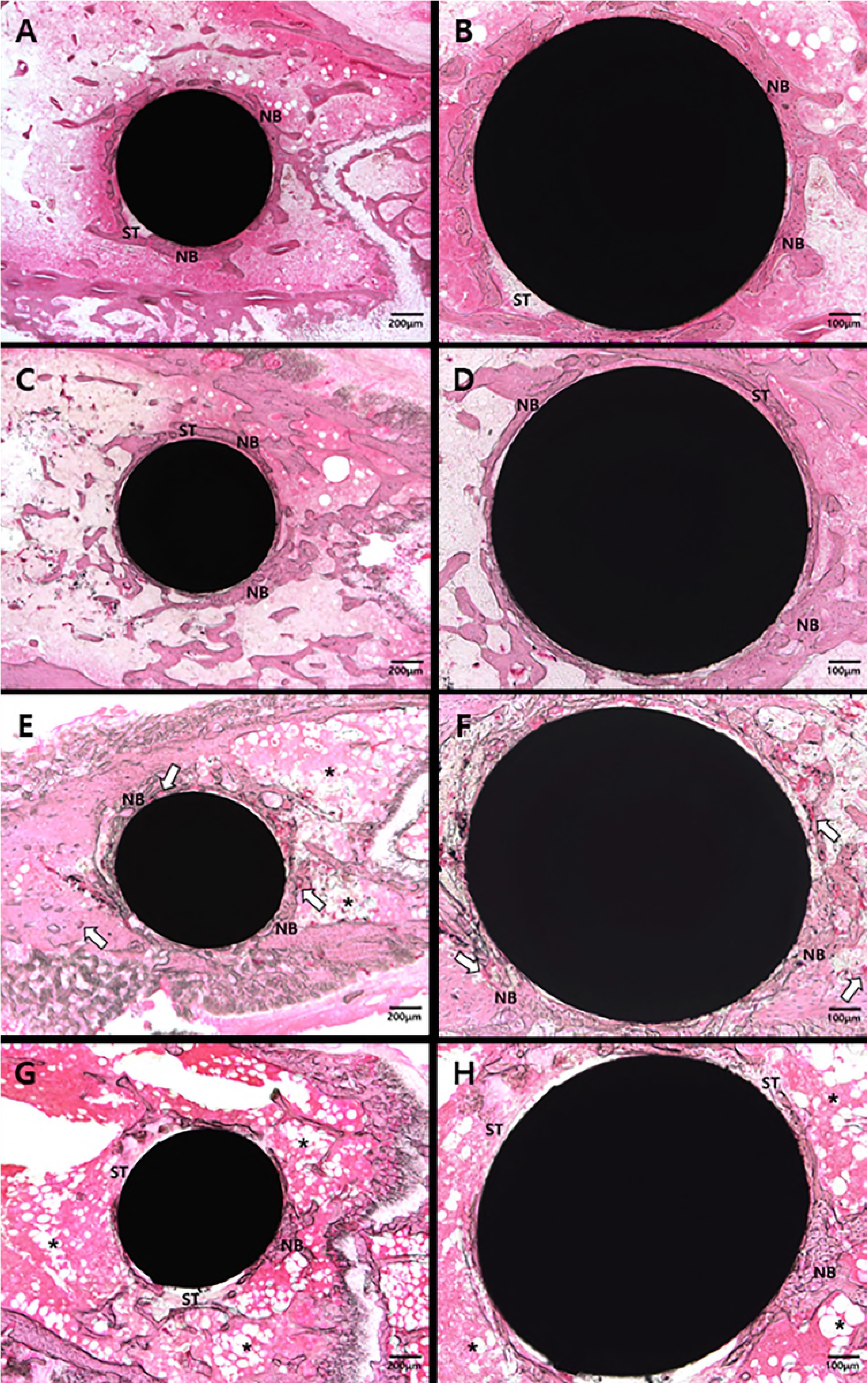
Representative images of undecalcified specimens of four groups. (A) and (B) Abundant and well-developed new bone (NB) formation observed around implant in the NH group, with some soft tissue (ST) engagement. (C) and (D) Thin and well-defined new bone formation observed in the NP group. (E) and (F) Plentiful vascular sinusoids with red blood cells (white arrow) and newly-formed bone surrounding implant in the DH group. (G) and (H) Fibrotic tissue and adipose tissue (asterisk) surrounding implant in the DP group.

### Histomorphometric and statistical analysis

Osseointegration was observed in all specimens of NH, NP, DH groups. One specimen of the DP group did not show osseointegration and was omitted from statistical analysis. All groups in this study demonstrated normality in parametric test (Shapiro-Wilk test). The BIC of HIF-1α treated groups was significantly higher than that of placebo groups in both normal and diabetic mice. There was no significant BIC difference between the NP and DH groups (Fig 4A). Among diabetic mice, the DH group showed significantly greater BV than the DP group while the groups of normal mice did not show any significant differences in BV. Only the DP group showed significantly low BV among the four groups (Fig 4B).

**Fig 4.**
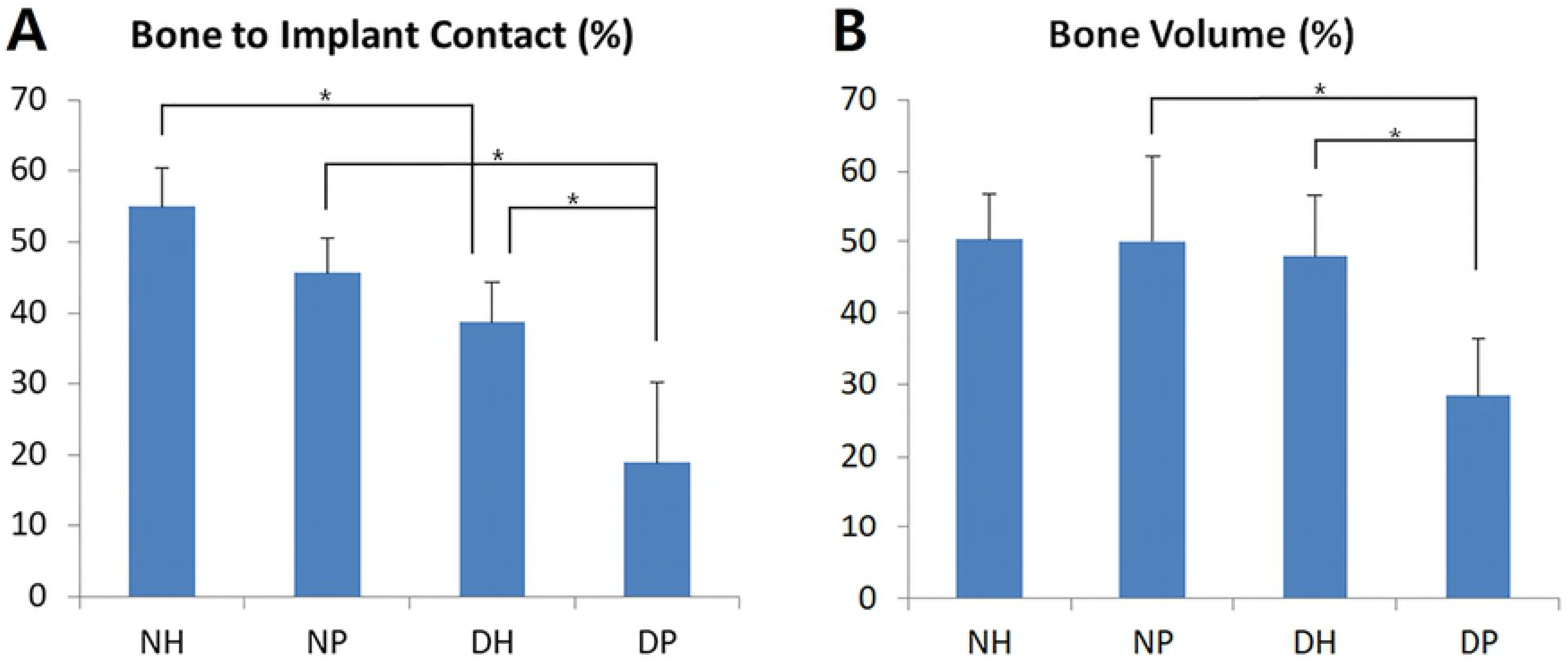
Histomorphometric analysis. (A) Linear percentage of direct BIC in the total surface of implants. (B) Percentage of newly formed bone area in circumferential zone within 100 um of the implant surface. Data represent mean **±** SD. *: p-value < 0.05, Normal mice with HIF-1α gel (NH); Normal mice with placebo gel (NP); Diabetic mice with HIF-1α gel (DH); Diabetic mice with placebo gel (DP).

## Discussion

Previous studies reported that hyperglycemia, even in hypoxic condition, negatively affects the stability and activation of HIF-1α and inhibits the expression of target genes of HIF-1α which are critical to wound healing [10]. On the other hand, it has been reported that the exogenous increase of HIF-1α resulted in improved bone regeneration and osseointegration around the implant [18,19]. This study was designed to test the hypothesis that the exogenous increase of HIF-1α would improve bone regeneration because endogenous HIF-1α was supressed in a hyperglycemic environment and failed to function based on these previous results.

According to Lu et al., immature mesenchymal tissue first appeared in significant amounts 4 days after bone ablation. Abundant formation of new bone was seen after 6 days. After 10 days, bone entered maturation stage and the bone marrow continuously repeated the remodeling cycle without major changes in the matrix [29]. Thus, specimens were obtained 2 weeks after the surgery to histologically evaluate mature bone in the healing process and estimate implant stability in each group.

In this study, histologic results show that adipose and soft tissue were more engaged in diabetic mice than in normal ones. Remarkably, most of the bone marrow in diabetic mice was composed of adipose tissue. In addition, the thickness and amount of bone were thinner and less in diabetic mice. Bone formation was highly irregular and fragile bone was also observed. The histological state of diabetic mice specimens was unfavorable for implant maintenance. These results were coincident with previous studies, which reported slower bone healing process in diabetic mice than normal mice as well as poorer biomechanical and histologic bone quality after initial healing [10,30].

Histomorphometric results showed that the DH group had greater bone contact and quantity than DP group. Based on histologic specimens, more vascular sinusoids were generated in those groups with HIF-1α application, demonstrating that HIF-1α increased the expression of VEGF and improved angiogenesis as in a previous study, in which HIF-1a was applied in the same way as this study [21]. It is speculated that HIF-1α, which enhanced angiogenesis, also enhanced bone regeneration and increased BIC and BV levels. In addition, distant sites were full of adipose tissue compared to HIF-1a application sites near the implant, which consisted mainly of dense bone. The difference was even more evident in comparison with the DP group, in which tissue around the implant was filled with adipose tissue.

The success of HIF-1α transferrence into the cell nucleus via PTD delivery system in this study is confirmed by RNA sequencing, which reveals 201 up-regulated genes and 15 down-regulated genes in the DH group compared with the DP group. Thus, one may conclude that HIF-1α had been delivered to the intracellular space and functioned as intended. Because factors related to bone formation were mostly expressed 4 days after implant surgery, RNA sequencing was performed at that time point [29].

21 target genes of HIF-1α out of 216 genes were selected through TRANSFAC^®^ and 6 genes related to tissue healing or bone regeneration in diabetes were found manually, based on previously reported research: *NOS2, GPNMB, CCL2, CCL5, CXCL16, TRIM63*. Functions of these genes are described in Table 5 [31–36].

**Table 5.**
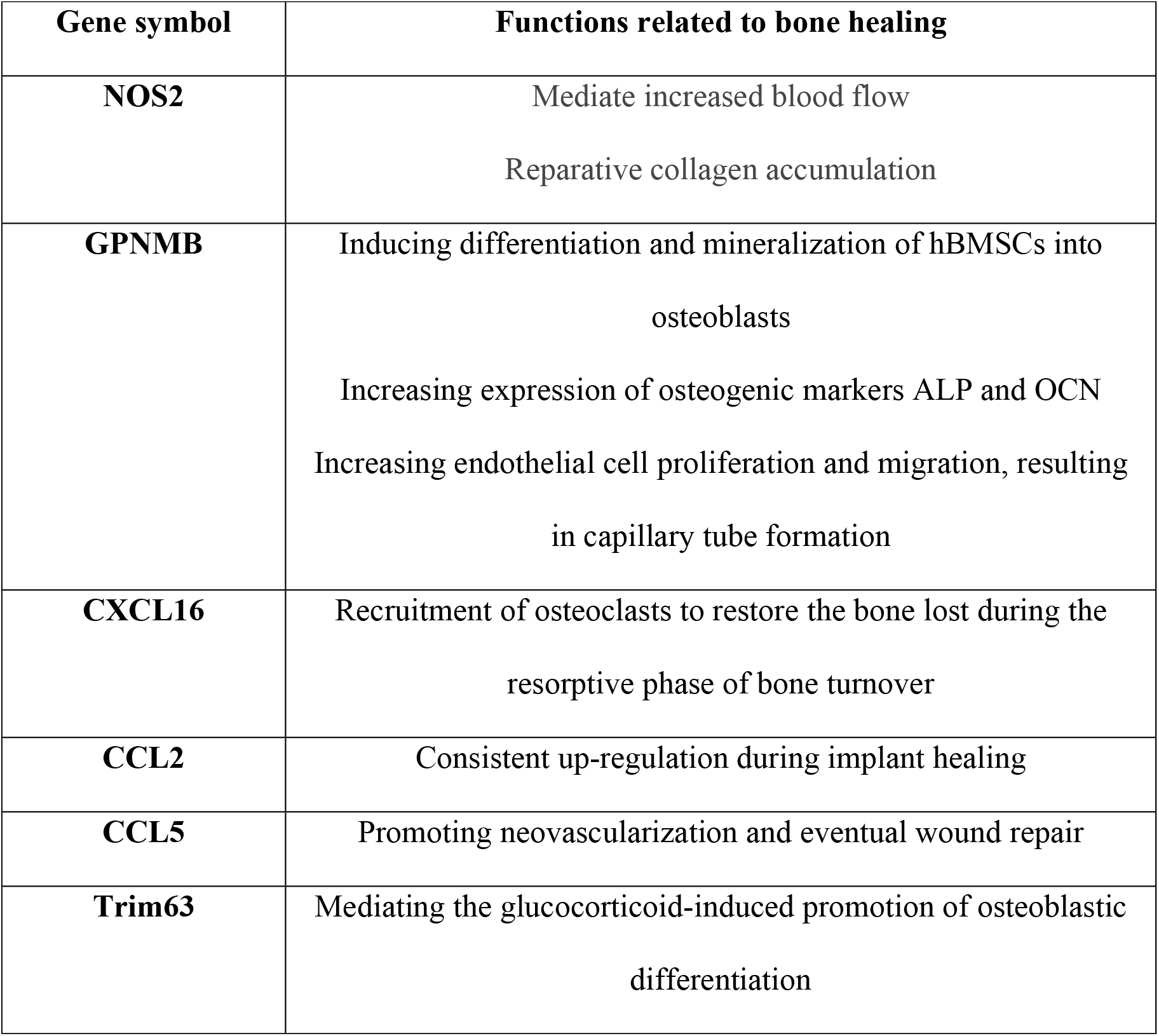
Genes related to tissue healing or bone regeneration selected out of 21 HIF-1α target genes in diabetic mice.

Bioinformatic analysis showed that diabetic mice had 216 DEGs and 21 target genes whereas normal mice had 95 DEGs and 5 target genes. Moreover, the DEGs and target genes in normal mice were mostly included in those of diabetic mice. For example, 3 genes out of 5 genes besides TNF and CD274 were also included as target genes within diabetic mice. These results suggest that application of HIF-1α, suppressed in a hyperglycemic condition, activated expression of HIF-1α target genes. On the other hand, HIF-1α target genes were less activated in a normal condition because endogenous HIF-1α was already sufficiently expressed around injured tissue. As a consequence, it is assumed that when exogenous HIF-1α was provided, genes downstream of HIF-1α were more actively expressed in diabetic mice, which may account for the histomorphometric results of improved BIC and BV

In normal conditions, HIF-1α also improved BIC and BV, consistent with previous studies [15,18]. However, HIF-1 a was more effective in diabetic mice in that the DH group showed significantly increased BIC and BV than DP while the NH group did not show a significant difference in BIC and BV compared to the NP group. Based on this result, it is assumed that exogenous HIF-1α worked more effectively in a hyperglycemic condition than normoglycemic.

Previous studies used mesenchymal stem cells to increase expression of HIF-1α, a process considered inefficient due to the long and complicated preparation [18,37]. On the other hand, the PTD-mediated DAN delivery system used in this study is a simple and efficient method for applying HIF-1α expression plasmid which can be easily produced in large quantities and can be injected [21]. Moreover, the molecular complex containing Hph-1-GAL4-DBD and HIF-1α-UAS can be solidified into gel form for application to a local site without diffusion. However, additional studies are needed to determine whether the gel form releases HIF-1α ideally or not compared to liquid type. Moreover, future studies on higher mammals such as dogs and pigs may clarify the effects of HIF-1α.

Bone regeneration can be enhanced by applying bone morphogenetic factor (BMP), an osteogenic growth factor, or vascular endothelial growth factor (VEGF), an angiogenic growth factor [38]. The administration of BMP has been reported to improve bone regeneration in numerous dental studies [39]. However, several complications associated with BMP treatment have been reported. Jeon demonstrated that BMP causes problems such as uncontrolled release rates, a short period of BMP release, and a high initial burst of release [40]. BMP also causes an unexpected immune reaction, spontaneous swelling of soft tissues, and difficulty in controlling diffusion [41]. HIF-1α is a transcription factor that promotes the expression of VEGF to induce angiogenesis and mediates callus remodeling and ossification through osteogenic differentiation [42,43]. VEGF expression by HIF-1α promotes angiogenesis by increasing the proliferation, survival, migration, and tube formation of endothelial cells [44,45]. Based on this study, we would like to suggest that the use of angiogenic growth factors such as HIF-1 α rather than osteogenic growth factors like BMP could improve bone quality and quantity around the implant.

## Conclusion

Regenerative strategies for diabetic patients in oral implantology require intact healing of localized bone defects. The success of the therapeutic procedure depends primarily on overcoming dysfunctional angiogenesis and restoring osteogenesis under hyperglycemic conditions. Innovative strategies include PTD-mediated HIF-1α gene delivery into the implanted sites locally to increase HIF-1α levels, giving rise to a hyperglycemic environment that favors regeneration. The implementation of gene-forced expression of HIF-1α delivered via PTD in such strategies holds tremendous potential. Further study will determine the potential effectiveness of a local delivery system with PTD.

## Acknowledgements

We thank DNA Link for helping conduct DNA extraction and analysis samples. Furthermore, we want to thank Se-young Kang and Yeun-Ju Kim for their assistance.

## Supporting information

**S1 File. Animal Research: Reporting of In Vivo Experiments (ARRIVE) guidelines.**

